# nsGSLs on tumors impair anti-tumor immune responses by OT-I T cells *in vitro* and support tumor growth *in vivo*

**DOI:** 10.1101/2024.10.13.617339

**Authors:** Tamara Verkerk, Aimée A.F. Selten, Nordin D. Zandhuis, Tao Zhang, Manfred Wuhrer, Robbert M. Spaapen, S. Marieke van Ham

## Abstract

Tumors often evolve to engage numerous strategies to circumvent detection by the immune system. Our group recently discovered elevated neolacto-series glycosphingolipids (nsGSL) surface levels as a possible immune evasion mechanism of tumors. We demonstrated a direct disruption of both innate and adaptive anti-tumor immunity *in vitro* when expression of nsGSLs was upregulated on established target cells. It remains unclear however, whether *in vivo* nsGSLs play an active role in tumor development and can aid tumors in evading immune responses.

To investigate whether nsGSLs facilitate tumor progression *in vivo*, we first established a murine model system using MC38-OVA cell lines with varying cell surface levels of nsGSLs. *In vitro* analysis revealed reduced MHC-I accessibility on tumor cells with elevated nsGSLs profiles, leading to diminished activation of OVA-specific OT-I T cells as evidenced by decreased expression of CD25, CD69, and production of IFNγ, which subsequently resulted in decreased tumor cell death. Subsequent *in vivo* experiments investigating tumor outgrowth after engraftment of subcutaneously injected MC38-OVA cell lines with low or high cell surface levels of nsGSLs demonstrated better growth of nsGSL-rich tumor cells compared to nsGSL-poor tumors which could be controlled. Together these results suggest that nsGSLs expressed by tumors can facilitate immune evasion and subsequent tumor progression. These data pave the way to explore whether targeting of the GSL pathway with specific inhibitors could be advantageous as a therapy against tumors with high nsGSL levels.

## Introduction

Tumors may exploit various mechanisms to modulate accessibility of important plasma membrane molecules to escape from recognition by immune cells. Well-studied is the alteration or inhibition in plasma membrane expression of the Major Histocompatibility Complex I (MHC-I, Human Leukocyte class I, HLA-I) which prevents presentation of (neo)antigens and subsequent recognition by tumor-specific cytotoxic T cells. Reversely, specific MHC-I alleles binding inhibitory receptors on immune cells can be upregulated to prevent their activation^1–8^. Other mechanisms include downregulation of immune activating proteins such as MHC-I polypeptide–related sequence A (MICA) or -B, which normally activates NK or γδ T cells through binding NKG2D^9,10^. Also upregulation of so-called immune checkpoint ligands such as PD-L1 (programmed death ligand 1) and CTLA4 (cytotoxic T-lymphocyte associated protein 4) is often used by tumor cells to avoid immune activation^9,11–16^. Recently, our group added variation in expression of the neolacto-series glycosphingolipids (nsGSL) to this list as a mechanism that is hijacked by tumors to assist escape from immune recognition and anti-tumor immune responses^17,18^.

Glycosphingolipids are membrane resident, glycan structures attached to a lipid base which are found on cells in varying compositions performing several functions which serve to maintain cellular homeostasis^19^. For example, they are involved in many cellular processes such as embryonic development, proliferation, apoptosis, adhesion, cell-cell or pathogen interactions protein function and signal transduction pathways^20,21^. All GSLs have a membrane-resident core consisting of a ceramide tail. Synthesis of the glycan on this tail is catalyzed by a series of enzymes in the Golgi system. First a glucose is added by the enzyme UGCG (UDP-glucose ceramide glucosyltransferase) and a galactose by B4GALT5/6 (β1,4-galactosyltransferase 5 or 6) which is used as a base to generate different GSL subtypes through addition of different sugars by different enzymes. For example, subsequent addition of a N-acetylglucosamine (GlcNAc) by B3GNT5 (β1,3-*N*-acetylglucosaminyltransferase 5) generates the group of neolacto-series GSLs (nsGSLs). Recently, we found that signal peptide protease like 3 (SPPL3) can control nsGSL synthesis through proteolytic destruction of B3GNT5. There are reports on aberrant levels of B3GNT5 expressed by tumors may point to a role of nsGSLs in tumor initiation, differentiation and/or progression. For example, Wang and colleagues found that bone marrow samples of patients with acute myeloid leukemia (AML) had a much higher expression of B3GNT5 than those of healthy individuals^22^. Similar observations for B3GNT5 expression were made for glioblastoma and, importantly, patients with high B3GNT5 expression had a shorter overall survival^23^.

Although aberrant GSL profiles are observed in several tumor types and correlations are found with disease progression, in these settings it is not clear whether nsGSLs actively contribute to tumor immune escape.

We previously demonstrated that aberrant expression of nsGSLs impair immune responses instigated by αβ T cells *in vitro*^17^. Activation and anti-tumor functionality of CD8+ T cells cultured with nsGSL-rich cells were diminished, as illustrated by decreased cytokine production and tumor cell killing respectively. CD8+ T cell function could be restored by knockout of the nsGSL producing enzyme B3GNT5. Interestingly, HLA-I on the cell surface of cells with high nsGSL levels was less accessible for antibodies and other proteins, which strongly implies that nsGSLs block the access of HLA-I for the TCR. Moreover, we showed in a sequential study that the accessibility of several other cell surface proteins on tumor cells is impaired for access by other immune cells. As a consequence, the anti-tumor cytotoxicity by innate immune cells such as γδ T cells and neutrophils towards nsGSL-rich tumor cells was hampered^18^.

Although we showed direct impairment of both innate and adaptive anti-tumor responses *in vitro*, it is yet unknown whether nsGSLs can support tumor development and play a role in their evasion of anti-tumor immune responses *in vivo*.

To uncover whether nsGSLs promote tumor progression *in vivo*, we used a murine model system entailing MC38 cell lines with either a low or high nsGSL content for evaluation of the effects of nsGSLs on tumor growth *in vivo*. We first established MC38 cell lines that express the well-known MHC-I binding model peptide SIINFEKL (OVA). After establishment of single cell (monoclonal) derived MC38-OVA cell lines, high nsGSL-expressing MC38-OVA cells were subsequently generated through consecutive knockout (KO) of murine Sppl3 (mSppl3) and overexpression of murine B3gnt5 (mB3gnt5). We showed that MHC-I accessibility was reduced due to overexpression these nsGSLs. In vitro experiments demonstrated that OVA specific OT-I T cell activation was diminished in response to the high nsGSL-expressing MC38-OVA, as measured by CD25, CD69 and IFN gamma (IFNγ) and yielded less tumor cell death of the high nsGSL-expressing MC38-OVA compared to the MC38-OVA control. Furthermore, an *in vivo* experiment showed prolific outgrowth of nsGSL-rich tumor cells while nsGSL poor tumors were better controlled. This indicates that nsGSLs expressed by tumors may promote tumor immune escape and subsequent tumor progression. Therefore, it may be beneficial to combine present immunotherapies with GSL inhibitors for the treatment of nsGSL rich tumors.

## Materials and Methods

### Mice

Female C57BL/6j/Ly5.1^+^Ly5.2^+^ and C57BL/6j/Ly5.2^+^ OT-I mice were bred and housed in the animal facility of the Netherlands Cancer Institute (NKI). All experiments were performed in accordance with institutional and national guidelines and permitted by the experimental animal committee of the NKI. Mice were included into the experiments at six to twelve weeks of age.

### Cell isolation and culture

#### Cell lines

MC38 cell lines (derived from C57BL/6, H-2k^b^/H-2D^b^, kindly gifted by Dr. Ramon Arens) were cultured in IMDM (Gibco) supplemented with 10% FCS (Serana) and antibiotics (Penicillin-Streptomycin, Invitrogen). HEK293T cells were cultured in DMEM supplemented with 10% FCS, 1% L-glutamine (Gibco) and 0.05 mM 2-mercapto-ethanol (bME (Sigma)). PlatE cells (kind gift by dr. Monika Wolkers) were cultured in DMEM (4.5 g/l D-glucose, L-glutamine and 25mM HEPES) supplemented with 10% FCS and antibiotics (Penicillin-Streptomycin). Primary OT-I T cells were purified from murine spleens, cultured and preactivated as described before^24^. All cells were cultured at 37°C and 5% CO_2_.

### Cell engineering

The primers and gRNAs used are listed in supporting tables I and II.

#### OVA expression

A pMXs-Puro vector (Cell Biolabs) was equipped with an RFP tag into the multiple cloning site^17^. The SIINFEKL sequence was ligated into the 5’end of the coding sequence of the puromycin gene in this pMXs-RFP plasmid (pMXs-RFP-SIINFEKL), destroying the puromycin gene (supporting table I-a). For virus production, the plasmid was transfected into Phoenix-Ampho cells (ATCC) supported by 11.5mM CaCL_2_ (Merck) and HBSP (50mM HEPES (Merck), 10mM KCl (Merck), 12 mM dextrose (Difco/Fisher Scientific), 280 mM NaCl (Merck) and 1.5mM Na_2_PO_4_ (pH 7.05, Merck)). After two full days of cultivation, the supernatant containing the retrovirus particles was harvested and filtered through a 0.2 μm filter. MC38-WT cells were transduced with the virus through spinoculation in presence of 8 μg/ml protamine sulfate (Merck). Transfection and transduction efficiencies were determined by RFP expression through flow cytometry. Next, cells were FACS sorted for RFP positive cells to obtain a pure population and, subsequently, monoclonal cell lines were generated through limiting dilution plating. SIINFEKL/OVA peptide expression of each clone was validated by activation of an OT-I hybridoma reporter cell line after coculture with the MC38-OVA clones. The best performing clone was selected for subsequent modification steps.

#### mSppl3 and mUgcg KO

gRNA targeting mouse Sppl3(mSppl3) or mouse Ugcg (mUgcg) were ligated into a lentiCRISPR-v2-GFP vector (Addgene) using BsmBI (New England Biolabs, supporting table I-b). The plasmid containing the gRNA targeting mSppl3 or mUgcg (lentiCRISPR-v2-m*Sppl3*gRNA-GFP or lentiCRISPR-v2-m*Ugcg*gRNA-GFP, Addgene) was co-transfected with packaging plasmids psPAX2, pVSVg and pAdVAntage (Promega) into HEK293T cells using polyethylenimine (PEI; Polyscience) for virus production. After two full days of cultivation, supernatant containing virus was filtered and stored at -80°C or used directly for transduction of MC38-OVA cells through spinoculation together with 8 μg/ml protamine sulfate. Transduced cells were (repetitively) FACS sorted for GFP positive cells until GFP expression remain stable >99% of the cells. The genetic knockout of mSppl3 or mUgcg was validated through deep amplicon sequencing by Genewiz (Azenta Life Sciences, supporting table II) combined with analysis using CRISPResso2 from the Pinellolab^25^.

#### mB3gnt5 overexpression

Mouse B3gnt5 was cloned into a pMX-puro plasmid (Cell Biolabs, supporting table I-a). For virus production, PlatE cells were transfected with either the pMX-puro-mB3gnt5 or an empty pMX-puro vector using the transfection reagent Genejammer (Agilent) according to the manufacturer’s protocol. After two full days of cultivation, the virus-containing supernatant was harvested and filtered through a 0.45 μm filter. The virus was directly used or stored at -80°C. MC38-OVA mSppl3^-/-^ cells were transduced with either the virus containing the pMX-puro-mB3gnt5 plasmid or the pMX-puro ‘empty vector control’ through spinoculation together with 8 μg/ml protamine sulfate. Next, the transduced cells were selected using puromycin (1 μg/ml).

### Glycosylation analysis

MC38-OVA cell lines were prepared for, and analyzed with porous graphitized carbon (PGC) LC-MS as described previously^17^.

### Cocultures

MC38-WT, MC38-OVA with low nsGSLs (mSppl3^-/-^EV^+^) and MC38-OVA with high nsGSLs (mSppl3^-/-^ mB3gnt5^+^) target cell lines were plated at 50,000 cells/well and left untouched for three hours to allow cells to adhere before coculture with OT-I cells in 100 μl IMDM (10% FCS and antibiotics. Next, OT-I cells were added to the MC38 cells at an effector to target (E:T) ratio of 0:1, 5:1 or 10:1 in a total volume of 150 μl. After a 24-hour coculture, cells were harvested and transferred to a V-bottom plate, after which the percentage of dead target cells (Near-IR positive, CD8 negative) and OT-I T cell (Near-IR negative, CD8 positive) activation through CD25 and CD69 expression were determined using flow cytometry.

For analysis of IFN gamma (IFNγ) producing primary OT-I T cells over multiple days, MC38 cell lines expressing different levels of nsGSLs were plated as described above. After 24 hours, the OT-I T cells in coculture were transferred to a new batch of the same MC38 cells (plated three hours in advance) and cultured for another 24 hours (48-time point). This was repeated for the 72-hour time point. Two hours before the end of each coculture, brefeldin A (BFA, BD Biosciences) was added at 1 μg/ml. The OT-I T cells were harvested after 6, 24, 48 and 72 hours and transferred to a V-bottom plate, after which the percentage of IFNγ positive cells was determined using flow cytometry.

### Flow cytometry

Prior to staining, cells were washed with PBS/0.1%BSA. MC38 cell lines were stained with antibodies targeting H-2Kb conjugated to APC (AF6-88.5.5.3, eBioscience) H-2Db in PerCP-eFluor710 (28-14-8, Invitrogen) together with LIVE/DEAD Near-IR (Invitrogen) diluted in PBS for 30 minutes on ice in the dark. Next, cells were washed twice with PBS/0.1% BSA and resuspended in PBS/0.1% BSA for acquisition and analysis.

For the cocultures, target cells and OT-I T cell were incubated with anti-CD8 in APC or in PerCPCy5.5 (53-6.7, Biolegend), anti-CD25 in AF488 (PC61, Biolegend), anti-CD69 in BUV737 (H1.2F3, BD Horizon) and LIVE/DEAD NEAR-IR for 30 minutes on ice and in the dark. Hereafter, cells were washed twice with PBS/0.1% BSA and resuspended in PBS/0.1% BSA for direct analysis or to be fixed for intracellular staining using the BD Cytofix/Cytoperm fixation and permeabilization kit (BD Bioscience) according to the manufacturer’s protocol. For intracellular staining, cells were permeabilized using permeabilization buffer and incubated with anti-IFNγ conjugated to PE (XMG1.2, Invitrogen) diluted in permeabilization buffer (BD Biosciences) for 30 minutes on ice in the dark. Subsequently, cells were washed twice and resuspended in PBS/0.1% BSA prior to analysis. Stained cells were analyzed or sorted on a BD flow cytometer (either LSR-II, Fortessa, FACSymphony, ARIA-II) and analyzed using FlowJo™ software version 10.9.0 (Ashland, OR: Becton, Dickinson and Company; 2023).

### *In vivo* tumor engraftment and monitoring

Mice were injected subcutaneously with 1 million MC38-OVA (n=4) control cells or MC38-OVA high nsGSL cells (n=4) in the right flank. One tumor in the control group did not engraft. Mice were randomly assigned to a group prior to tumor cell injection and housed while being mixed. Tumor growth was measured on day 4, 8, 11, 14, 18, 22 and 26 after tumor engraftment in cubic millimeter (mm^3^). After 26 days, mice were sacrificed.

### Statistical Analysis

Statistical testing was done by a student’s T-test or a one-way ANOVA followed by a Tukey’s multiple comparison test. The statistical analysis was performed using GraphPad Prism version 10.0 for Mac OS (GraphPad Software, Boston, Massachusetts USA). Differences were considered significant when p≤0.05.

## Results

### nsGSLs affect the anti-tumor cell cytotoxicity of OT-I T cells

Previously, we demonstrated that the loss of SPPL3 in tumor target cells reduced the cytokine production of human CD8 T cells due to nsGSL mediated interference. To assess whether elevated nsGSL levels would have a similar effect in a murine system, we generated mouse tumor MC38 OVA cells with low and high nsGSL levels. The MC38 tumor cell line was chosen because it has been shown to be highly immunogenic in several in *vivo* studies^26–28^. Therefore, if nsGSL overexpression in this cell line may benefit anti-tumor immune escape, this may provide a window to analyse the effect of nsGSLs on anti-tumor immune responses.

First, we made a monoclonal OVA (SIINFEKL antigen) expressing MC38 cell line to ensure homogeneous OVA expression before generation of subsequent cell lines. Next, this cell line was knocked out of mouse Sppl3 (mSppl3^-/-^) or mouse Ugcg (mUgcg^-/-^) as a control for nsGSL profile analysis. To maximally promote nsGSL expression, mouse B3gnt5 (mB3gnt5^+^) was overexpressed in the mSppl3^-/-^ cells. Simultaneously, MC38-OVA mSppl3^-/-^ cells with an ‘empty vector’ (mSppl3^-/-^EV^+^) was created as a control cell line.

To validate nsGSL overexpression in the mSppl3^-/-^mB3gnt5^+^ cells, mass spectrometry of GSLs was used to analyse the GSL expression profiles on the generated cell lines. The mSppl3^-/-^ and mSppl3^-/-^EV^+^ cells showed a minimal elevation of nsGSLs compared to MC38-OVA WT, but overexpression of mB3gnt5 (mSPPL3^-/-^mB3gnt5^+^) resulted in high nsGSL levels (Figure 1). Cells lacking mUgcg contained no GSLs which was in line with the expectation.

**Figure 1.**
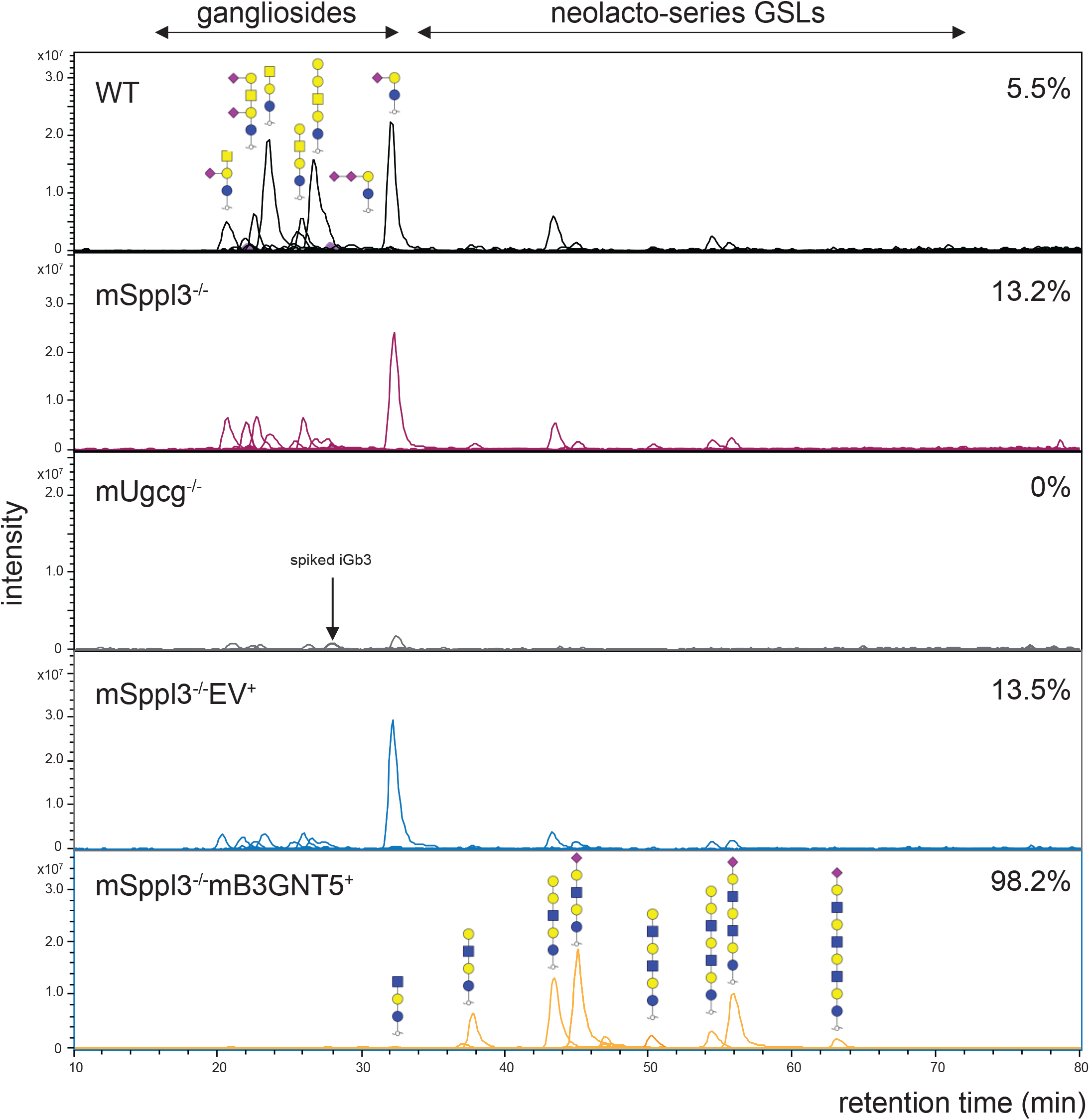
Chromatograms of PGC LC-MS on total GSL glycans. Extracted ion chromatograms of total GSL glycan content released from MC38-OVA ‘WT’, mSppl3^-/-^. mUgcg^-/-^, mSppl3^-/-^EV^+^ and mSppl3^-/-^mB3gnt5^+^ cell lines using PGC LC-MS.

Next, we assessed whether the upregulated nsGSL levels impair the binding of antibodies targeting murine MHC-I similar to previous observations with human HLA-I^17^. We stained MC38-OVA mSppl3^-/-^ev^+^ and mSppl3^-/-^B3gnt5^+^ cells for H-2K^b^ and H-2D^b^ expressed by C57BL/6 mice^29^. Both H-2K^b^ and H-2D^b^ cell surface staining of MC38-OVA cells with high nsGSL levels (mSppl3^-/-^B3gnt5^+^) was reduced compared to the control cells (mSppl3^-/-^EV^+^), indeed indicating a similar interference with MHC-I targeting antibodies as previously observed for human cells (Figure 2A). Next, it was evaluated whether nsGSLs have an effect on activation and tumor cell killing by OVA peptide (SIINFEKL) specific OT-I T cells. OT-I T cells were cocultured together with MC38-WT, MC38-OVA mSppl3^-/-^EV^+^ (control) and mSppl3^-/-^B3gnt5^+^ cells (high nsGSLs) in different ratios. At different effector to target ratios OT-I cells demonstrated a higher killing activity against the OVA antigen expressing control cells compared to the cells containing high nsGSL content (Figure 2B, C). Moreover, OT-I cells were less activated after coculture with MC38-OVA cells with high nsGSL levels, as evidenced by a decreased expression of the activation markers CD25 (10:1 E:T ratio) and CD69 (5:1 and 10:1 ratios) (Figure 2D, E).

**Figure 2.**
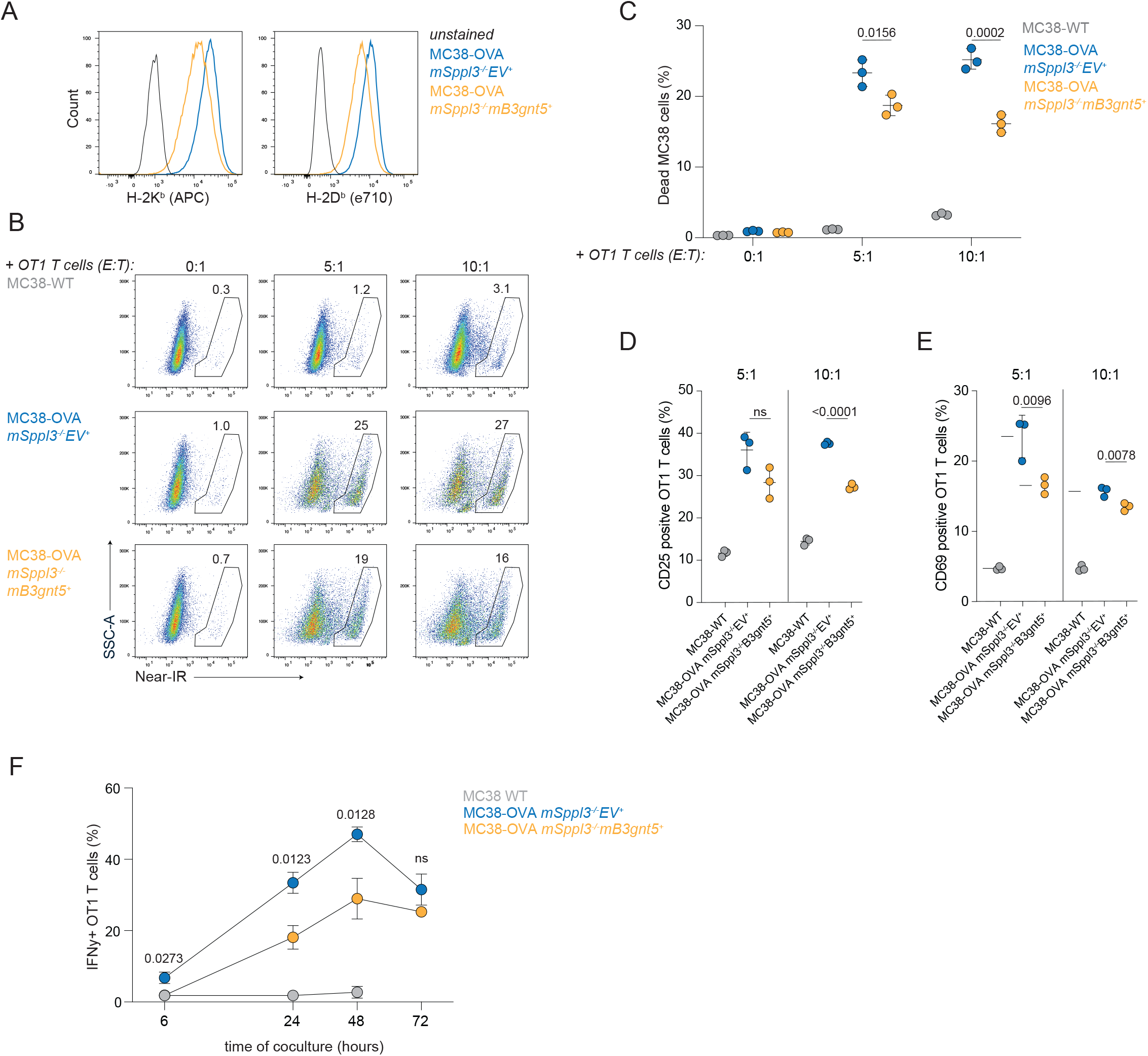
nsGSLs impair anti-tumor cytotoxicity of OT-I T cells. Freshly activated OT-I T cells were cocultured with MC38-WT or MC38-OVA cell lines containing low nsGSL levels (mSppl3^-/-^EV^+^) or high nsGSL levels (mSppl3^-/-^mB3gnt5^+^) for 24 hours or multiple days in different effector to target (E:T) ratios. **(A)** Histograms of MC38-OVA control cells (mSppl3^-/-^EV^+^) or with high nsGSL levels (mSPPL3^-/-^mB3GNT5^+^) stained for H-2K^b^ and H-2D^b^. Representative flow cytometry plots **(B)** and combined data of **(C)** showing the percentage of dead (Near-IR positive) MC38-WT, MC38-OVA mSppl3^-/-^EV^+^ or mSppl3^-/-^mB3gnt5^+^ target cells with different E:T ratios. **(D, E)** The MFI of CD25 and CD69 expression by OT-I T cells. Each dot represents the average of triplicates of one OT-I batch (n=3). **(F)** The percentage of IFNγ positive OT-I T cells during after 6, 24, 48 and 72 hours of coculture. Each dot represents the average of three OT-I batches for which triplicates were used (n=3). An unpaired one-way ANOVA was used to assess statistical significances.

Another coculture was performed to investigate the effect of nsGSLs on functional OT-I T cell over time. Already after six hours, production of the inflammatory cytokine interferon gamma (IFNγ) by OT-I T cells cocultured with mSppl3^-/-^B3gnt5^+^ cells was reduced compared to the coculture with mSppl3^-/-^EV^+^ cells (Figure 2F). This reduction continued up to 2 days (48 hours) of coculture after which the IFNγ production of OT-I T cells decreased under both conditions.

Together, these data show that excessive nsGSL expression protects tumor cells from elimination by OT-I T cells through inhibition of OT-I T cell activation.

### Elevated nsGSL levels enhance tumor outgrowth *in vivo*

Given that nsGSLs protect tumor cells against anti-tumor immune responses *in vitro*, we then investigated to which extent nsGSLs support/drive escape from anti-tumor immune responses *in vivo*. Mice were injected with either MC38-OVA mSppl3^-/-^EV^+^ cells (low nsGSLs) or mSppl3^-/-^B3gnt5^+^ cells (high nsGSLs) and tumor growth was monitored for 26 days after which the animals were sacrificed. Tumors from both groups grew similar up to 14 days after tumor injection (Figure 3A). After 14 days, MC38-OVA mSppl3^-/-^ B3gnt5^+^ showed a robust growth while the MC38-OVA control tumors remained stable in size and one tumor disappeared completely (Figure 3A, B). Thus, nsGSLs promote tumor outgrowth *in vivo*.

**Figure 3.**
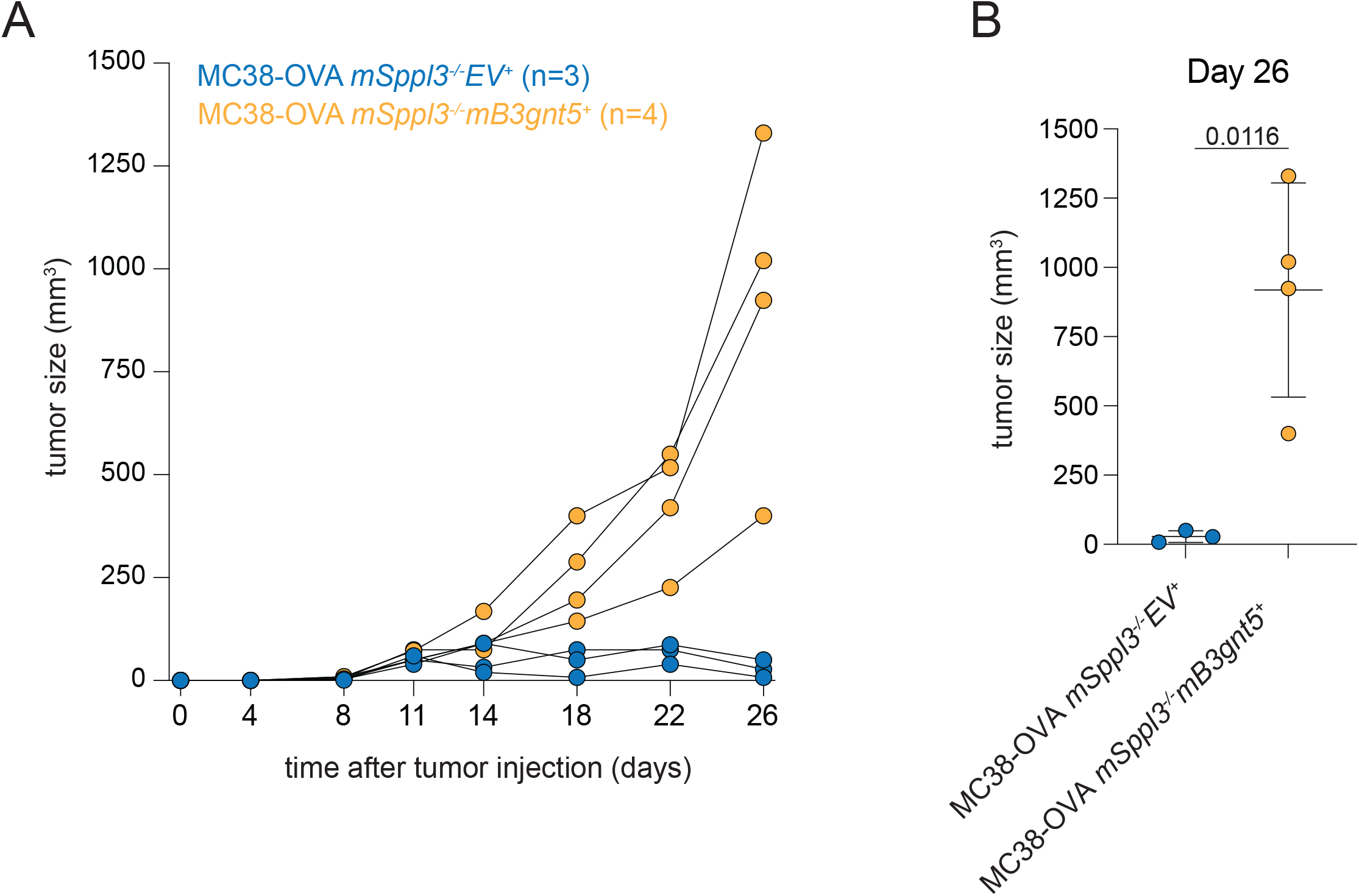
Tumors with high nsGSL profile grow prosperously while nsGSL low tumors can be controlled. Mice were injected with MC38-OVA cell lines containing low nsGSL levels (mSppl3^-/-^EV^+^, n=4) or high nsGSL levels (mSppl3^-/-^mB3nt5^+^, n=4) and tumor growth was monitored for 26 days. One tumor of the first group did not engraft. The tumor size (in mm^3^) of each group on each day of measurement is shown in **(A)** and a summary of the tumor size on day 26 is shown in **(B)**. A student’s T-test was used to assess statistical differences.

## Discussion

In this study, we showed that tumor cells expressing high amounts of nsGSLs are better protected against anti-tumor responses from tumor specific OT-I T cells *in vitro* as illustrated by a reduced percentage of dead tumor cells in comparison with low nsGSL expressing counterparts. In addition, the activation of the OT-I cells was reduced as demonstrated by lower expression of activation markers CD25 and CD69 and decreased IFNγ peak production. More importantly, tumors abundantly expressing nsGSLs showed robust outgrowth while nsGSL low tumors did not thrive. These data indicate for the first time that nsGSLs facilitate escape from anti-tumor immune responses *in vivo*.

MC38 cells naturally do not express many nsGSLs and thus it could be that in these cells either the Sppl3 gene is very active, that B3gnt5 gene is less active or a combination of both. Surprisingly, sole knockout of mSppl3 did not elevate the levels nsGSL as our group previously observed for human HAP1 cells^17^. Extra expression of mB3gnt5 strongly increased nsGSL levels as demonstrated with mass spectrometric analysis, supporting the theory of low B3gnt5 activity in unmodified MC38 cells. We did not test the sole overexpression of mB3gnt5. To validate low mB3gnt5 further, nsGSL profiles of single mB3gnt5 overexpressing MC38 cells should be analysed in future experiments.

The *in vivo* experiment that was performed in this study showed vigorous outgrowth of tumors with excessive nsGSL expression. The tumors in the control group initially engrafted, but failed to progress. A potential explanation for this difference is reduced recognition of nsGSL high MC38 tumors by CD8+ T cells compared to tumors with low nsGSL expression. This is in line with the *in vitro* results in figure 2, where we show impaired OT-I activation and MC38-OVA cell killing in the situation of elevated nsGSL profiles compared to cells with low nsGSL profiles. Moreover, it is likely that this is driven by the hampered accessibility of MHC-I as we observed a lower cell surface antibody staining for MHC-I (H-2K^b^ and H-2D^b^). In this *in vivo* study, however, we did not investigate the tumor infiltration CD8+ T cell population or their antigen specificity. Therefore, we do not know if there is a difference in the occurrence of tumor antigen specific CD8+ T cells in nsGSL-rich or nsGSL-poor MC38-OVA tumors.

Comparing the occurrence and specificity of tumor infiltrating T cells between nsGSL high and low tumors may provide more insights into the role of nsGSLs MC38-OVA tumor cells recognition. Like the OT-I T cells used *in vitro*, the endogenous T cells could be OVA-specific, which has been shown for endogenous MC38-OVA tumor infiltrating CD8+ T cells by others^30^. In addition to OVA antigen presentation, these cell lines may have also presented other immunogenic antigens. For example, we introduced GFP in these cells as a selection marker for MC38-OVA mSppl3^-/-^ cells and peptides derived from GFP could potentially be presented by H-2K^b^ and/or H-2D^b^. By several studies, expression of GFP has been shown to be immunogenic^31,32^. For example, Grzelak and colleagues showed that endogenous GFP-specific CD8+ T cells were present in BALB/c mice (H-2D^b^/H-2D^d^) engrafted with 4T1-GFP+ tumor cells using H-2D^d^ (MHC-I) tetramers^32^. To our knowledge, it has not been shown that GFP derived peptides can also be presented by H-2K^b^ and/or H-2D^b^ which are the MHC-I alleles expressed by MC38 cells. Therefore, it should be further investigated whether GFP derived antigens are indeed presented by the MC38-OVA cells and could induce a CD8+ T cell mediated immune response.

An additional explanation for the reduced growth of nsGSL-poor tumors could be that the expression levels of either OVA or GFP is higher compared to nsGSL-rich tumors, which renders these cells more immunogenic. This is, however, unlikely because the nsGSL-rich and nsGSL-poor MC38-OVA cell lines originate from the same monoclonal MC38-OVA cell line. In addition, both cell lines were sorted for the same GFP intensity indicating equal GFP expression in these MC38-OVA cell lines.

To our knowledge, our data provide the first *in vivo* evidence of impaired tumor control due to overexpression of specifically nsGSLs. Several studies identified human tumor types containing nsGSL-skewed profiles compared to healthy equivalents, for example in AML^22^. In addition, B3GNT5 activity has been correlated with tumor progression and metastasis^23,33^. Yet, the specific role for B3GNT5 and nsGSLs in tumor progression was not clear. Here we make the comparison between nsGSL-poor and nsGSL-rich tumors directly and found that nsGSLs can support tumor growth.

Nonetheless, we recognize that the *in vivo* experiment demonstrated here was performed once and involved a limited number of mice. Thus, this observation requires further validation. Still, our group has demonstrated through a number of studies that nsGSLs impair immune cell activation, cytokine production and tumor cell killing *in vitro*^17,18^. Combining the information gathered *in vitro and in vivo*, there is a strong suggestion that nsGSLs play a notable role in escape from anti-tumor immunity.

Future *in vivo* experiments should include phenotyping of the tumors *ex vivo* to assess the composition of endogenous tumor infiltrating lymphocytes and their tumor specificity and activation and differentiation status. This could provide insight in the development and interplay of different immune cells, such as T cells, B cells, NK cells and γδ T cells, in the anti-tumor immune response against high nsGSL containing tumors. Moreover, through adoptive cell transfer of OT-I T cells, combined with phenotyping of activation and exhaustion markers, one may gain more knowledge on the specific anti-tumor response and differentiation of already activated T cells.

Here, we presented a role for nsGSLs in *in vivo* tumor growth in addition to new and previously generated *in vitro* data indicating a pivotal role for targeting these membrane-bound structures in the fight of immune cells against tumors. Because the GSL synthesis pathway is safely inhibited in lysosomal storage diseases, our combined data warrant investigations on the efficacy of GSL synthesis inhibition to treat patients with nsGSL-rich tumors.

## Acknowledgements

We thank the animal caretakers of the NKI for their support and we would like to thank the Sanquin Research core facility and colleagues from Sanquin; M. Hoogenboezem, A.R.G Laurent, N.A.M. Kragten, H. Heimans, S. J. Bliss, T. Jorritsma and K.P.J.M. van Gisbergen. This research was supported by grants from the Dutch Research Council (Grant NWO-VIDI 91719369 to RS) and the Landsteiner Foundation for Blood Transfusion Research (LSBR Fellowship 1842F to RS) and an internal Product and Process Development grant (PPOC) of Sanquin with number L2593.

---

**Supporting table I-a.**
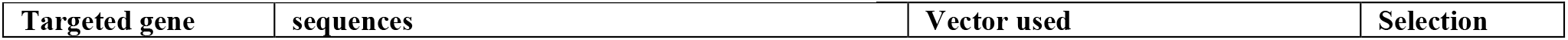
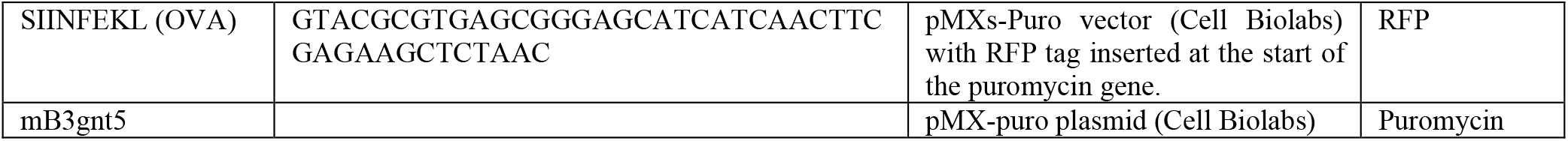
Details on (over)expressed genes.

**Supporting table I-b.**
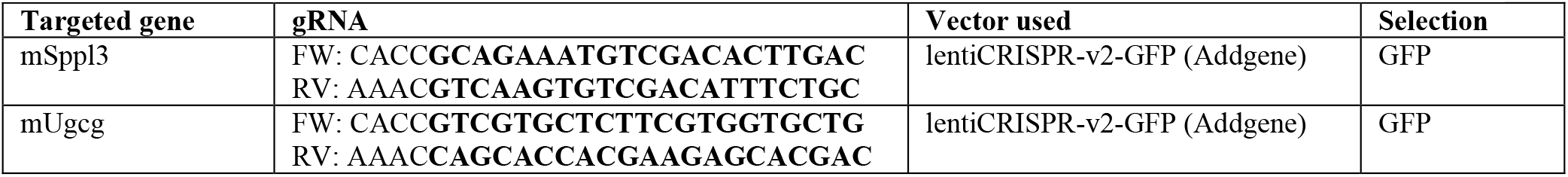
Details of CRISPR/CAS mediated genome editing. gRNA: BsmBI cutting-site compatible overhang followed the gRNA (bold).

**Supporting table II.**
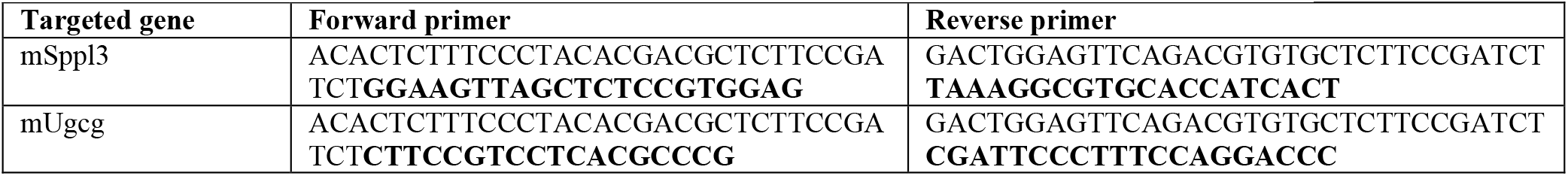
Primers used for deep sequencing. Primers: amplicon sequence directly followed by the primer sequence (bold).

## References

1. Dhatchinamoorthy, K., Colbert, J. D. & Rock, K. L. Cancer Immune Evasion Through Loss of MHC Class I Antigen Presentation. Front. Immunol. 12, (2021).

2. Liu, L., Wang, L., Zhao, L., He, C. & Wang, G. The Role of HLA-G in Tumor Escape: Manipulating the Phenotype and Function of Immune Cells. Front. Oncol. 10, 1–11 (2020).

3. Borst, L., van der Burg, S. H. & van Hall, T. The NKG2A-HLA-E axis as a novel checkpoint in the tumor microenvironment. Clin. Cancer Res. 26, 5549–5556 (2021).

4. Seliger, B. et al. HLA-E expression and its clinical relevance in human renal cell carcinoma. Oncotarget 7, 67360–67372 (2016).

5. Paschen, A. et al. Complete loss of HLA class I antigen expression on melanoma cells: A result of successive mutational events. Int. J. Cancer 103, 759–767 (2003).

6. Martínez-Jiménez, F. et al. Genetic immune escape landscape in primary and metastatic cancer. Nat. Genet. 55, 820–831 (2023).

7. Fangazio, M. et al. Genetic mechanisms of HLA-I loss and immune escape in diffuse large B cell lymphoma. Proc. Natl. Acad. Sci. U. S. A. 118, (2021).

8. McGranahan, N. et al. Allele-Specific HLA Loss and Immune Escape in Lung Cancer Evolution. Cell 171, 1259–1271.e11 (2017).

9. Xing, S. & Ferrari de Andrade, L. NKG2D and MICA/B shedding: a ‘tag game’ between NK cells and malignant cells. Clin. Transl. Immunol. 9, 1–10 (2020).

10. Schmiedel, D. & Mandelboim, O. NKG2D ligands-critical targets for cancer immune escape and therapy. Front. Immunol. 9, 1–10 (2018).

11. Raulet, D. H., Gasser, S., Gowen, B. G., Deng, W. & Jung, H. Regulation of ligands for the NKG2D activating receptor. Annual Review of Immunology vol. 31 (2013).

12. Cha, J. H., Chan, L. C., Li, C. W., Hsu, J. L. & Hung, M. C. Mechanisms Controlling PD-L1 Expression in Cancer. Mol. Cell 76, 359–370 (2019).

13. Vathiotis, I. A., Gomatou, G., Stravopodis, D. J. & Syrigos, N. Programmed death-ligand 1 as a regulator of tumor progression and metastasis. Int. J. Mol. Sci. 22, 1–13 (2021).

14. Rotte, A. Combination of CTLA-4 and PD-1 blockers for treatment of cancer. J. Exp. Clin. Cancer Res. 38, 1–12 (2019).

15. Pardoll, D. M. The blockade of immune checkpoints in cancer immunotherapy. Nat. Rev. Cancer 12, 252–264 (2012).

16. Miller, M. A., Sullivan, R. J. & Lauffenburger, D. A. Molecular pathways: Receptor ectodomain shedding in treatment, resistance, and monitoring of cancer. Clin. Cancer Res. 23, 623–629 (2017).

17. Jongsma, M. L. M. et al. The SPPL3-Defined Glycosphingolipid Repertoire Orchestrates HLA Class I-Mediated Immune Responses. Immunity 54, 132–150.e9 (2021).

18. Verkerk, T., Waard A. A. De, Koomen, S. J. I. & Sanders, J. Tumor-expressed SPPL3 supports innate anti-tumor immune responses. (2024).

19. Merrill, A. H. Sphingolipid and glycosphingolipid metabolic pathways in the era of sphingolipidomics. Chem. Rev. 111, 6387–6422 (2011).

20. Zhang, T., De Waard, A. A., Wuhrer, M. & Spaapen, R. M. The role of glycosphingolipids in immune cell functions. Front. Immunol. 10, 1–22 (2019).

21. D’Angelo, G., Capasso, S., Sticco, L. & Russo, D. Glycosphingolipids: Synthesis and functions. FEBS J. 280, 6338–6353 (2013).

22. Wang, Z. et al. High expression of lactotriaosylceramide, a differentiation-associated glycosphingolipid, in the bone marrow of acute myeloid leukemia patients. Glycobiology 22, 930–938 (2012).

23. Jeong, H. Y. et al. B3GNT5 is a novel marker correlated with stem-like phenotype and poor clinical outcome in human gliomas. CNS Neurosci. Ther. 26, 1147–1154 (2020).

24. Zandhuis, N. D. et al. Regulation of IFN γ production by ZFP36L2 in T cells is. (2024).

25. Clement, K. et al. CRISPResso2 provides accurate and rapid genome editing sequence analysis. Nat. Biotechnol. 37, 224–226 (2019).

26. Schrörs, B. et al. MC38 colorectal tumor cell lines from two different sources display substantial differences in transcriptome, mutanome and neoantigen expression. Front. Immunol. 14, 1–10 (2023).

27. Juneja, V. R. et al. PD-L1 on tumor cells is sufficient for immune evasion in immunogenic tumors and inhibits CD8 T cell cytotoxicity. J. Exp. Med. 214, 895–904 (2017).

28. Greenlee, J. D. & King, M. R. A syngeneic MC38 orthotopic mouse model of colorectal cancer metastasis. Biol. Methods Protoc. 7, (2022).

29. Moussa, P., Marton, J., Vidal, S. M. & Fodil-Cornu, N. Genetic dissection of NK cell responses. Front. Immunol. 3, (2012).

30. Kurtulus, S. et al. Checkpoint Blockade Immunotherapy Induces Dynamic Changes in PD-1 − CD8 + Tumor-Infiltrating T Cells. Immunity 50, 181–194.e6 (2019).

31. Schultheiß, C. & Binder, M. Overcoming unintended immunogenicity in immunocompetent mouse models of metastasis: the case of GFP. Signal Transduct. Target. Ther. 7, 3–4 (2022).

32. Grzelak, C. A. et al. Elimination of fluorescent protein immunogenicity permits modeling of metastasis in immune-competent settings. Cancer Cell 40, 1–2 (2022).

33. Meng, Q. et al. A circular network of coregulated sphingolipids dictates lung cancer growth and progression. EBioMedicine 66, (2021).

